# PHOSPHORYLATION OF RyR2 SIMULTANEOUSLY EXPANDS THE DYAD AND REARRANGES THE TETRAMERS

**DOI:** 10.1101/2023.05.23.541024

**Authors:** Parisa Asghari, David R.L. Scriven, Saba Shahrasebi, Hector H. Valdivia, Xander H.T. Wehrens, Edwin D.W. Moore

## Abstract

We have previously demonstrated that type II ryanodine receptors (RyR2) tetramers can be rapidly rearranged in response to a phosphorylation cocktail. The cocktail modified downstream targets indiscriminately making it impossible to determine whether phosphorylation of RyR2 was an essential element of the response. We therefore used the β-agonist isoproterenol and mice with one of the homozygous mutations, S2030A^+/+^, S2808A^+/+^, S2814A^+/+^, or S2814D^+/+^, to address this question and to elucidate the role of these clinically relevant mutations. We measured the length of the dyad using transmission electron microscopy (TEM) and directly visualized RyR2 distribution using dual-tilt electron tomography. We found that: 1) The S2814D mutation, by itself, significantly expanded the dyad and reorganized the tetramers suggesting a direct link between the phosphorylation state of the tetramer and the microarchitecture. 2) All of the wild-type, as well as the S2808A and S2814A mice, had significant expansions of their dyads in response to ISO, while S2030A did not. 3) In agreement with functional data from the same mutants, S2030 and S2808 were necessary for a complete β-adrenergic response, whereas S2814 was not. 4) All the mutated residues had unique effects on the organization of their tetramer arrays. 5) The correlation of structure with function suggests that tetramer-tetramer contacts play an important functional role. We conclude that both the size of the dyad and the arrangement of the tetramers are linked to the state of the channel tetramer and can be dynamically altered by a β-adrenergic receptor agonist.

**Summary:** Analysis of RyR2 mutants suggests a direct link between the phosphorylation state of the channel tetramer and the microarchitecture of the dyad. All phosphorylation site mutations produced significant and unique effects on the structure of the dyad and its response to isoproterenol.

## Introduction

The type II ryanodine receptor (RyR2) is a large, 2.2 MDa, homotetrameric Ca^2+^-activated Ca^2+^ ion channel located in the endoplasmic and sarcoplasmic reticulum of many cells(Lanner et al., 2010; Van Petegem, 2015; Meissner, 2017). In the heart, its distribution appears to be limited to the junctional sarcoplasmic reticulum (jSR), which in ventricular cardiomyocytes is located ∼ 15 nm distant from the sarcolemma and the transverse and axial t-tubules(Asghari et al., 2009). The close apposition of the adjacent membranes creates a space of restricted diffusion, the dyad, enabling the small current through Ca_v_1.2 to activate the underlying RyR2 through Ca^2+^- induced Ca^2+^ release (CICR) despite the tetramers’ relatively low affinity for Ca^2+^ (Carl et al., 1995; Sun et al., 1995; Franzini-Armstrong, 2018). Each cell has thousands of dyads (Scriven et al., 2010), and in systole the action potential coordinates their near simultaneous release of Ca^2+^ into the myoplasm to produce a coordinated contraction.

In diastole, tetramers open stochastically, but with low probability(Cheng et al., 1993; Satoh et al., 1997). A single tetramer opening may result in an isolated event, a Ca^2+^ quark (Lipp and Niggli, 1996), but since tetramers can be distributed in clusters, some with several hundred members (Baddeley et al., 2009; Hayashi et al., 2009), inter-RyR2 CICR can result in larger Ca^2+^ release events, blips (Iaparov et al., 2021) or sparks (Cheng et al., 1993; Lopez-Lopez et al., 1994), depending on the number that are opened. Diastolic Ca^2+^ release, or leak, is normal but becomes dangerously elevated in many inherited and acquired cardiac diseases, such as catecholaminergic polymorphic ventricular tachycardia (CPVT) and heart failure (Keefe et al., 2023). This has led to increased morbidity and mortality (Bers, 2014) which is driving efforts to understand how RyR2 is regulated.

Intuitively, the microarchitecture of the dyad should affect the spatial and temporal characteristics of Ca^2+^ sparks and diastolic Ca^2+^ release, and this has been demonstrated by mathematical models which have highlighted the importance of the number of tetramers in a cluster, their relative positions, density and coupling efficiency (Tanskanen et al., 2007; Cannell et al., 2013; Iaparov et al., 2019; Iaparov et al., 2021).

Direct examination of the tetramers’ positions in intact rat and human myocyte dyads using electron tomography has shown that the array is irregular but neither random nor static (Asghari et al., 2014). At rest, most of the tetramers are in contact with at least one other, in positions that were broadly characterized as side-by-side or checkerboard and in roughly comparable proportions. Factors that decreased the Ca^2+^ spark frequency, 4 mM Mg^2+^ and the immunophilins FKBP12 and FKBP12.6, produced clusters with significantly more tetramers in the side-by-side configuration at the expense of those in a checkerboard while those that increased the Ca^2+^ spark frequency, 0.1 mM Mg^2+^ and phosphorylation, did the opposite (Asghari et al., 2014; Asghari et al., 2020). Under all experimental conditions there were a few tetramers whose sides were not parallel to each other or were separated from their neighbours, and were classified as isolated.

The correlated structure and function suggested a simple model in which the receptor’s relative positions modifies their gating parameters in some manner, with changes in the ratio of the checkerboard to side by side predicting changes in the Ca^2+^ spark frequency. An increase in the ratio is associated with an increase in spark frequency while the converse is also true. The simplest hypothesis, given experimental evidence (Marx et al., 2001; Porta et al., 2012), is that the increase in spark frequency in the checkerboard configuration might be due to positive allosteric interaction. Negative allosteric interaction in the side-by-side configuration is also possible, but lacks experimental evidence. In this model, the mixed organization of checkerboard and side-by-side tetramers observed at rest in the fixed cells is a snapshot of a dynamic environment where the tetramers’ relative positions are in motion, shifting with changes in their phosphorylation status, bound ions and small molecules.

Our previous observations used permeabilized rat ventricular myocytes and a phosphorylation cocktail that activated numerous kinases while inhibiting phosphatases PP1 and PP2A (Asghari et al., 2014). This method produced indiscriminate phosphorylation of downstream targets making it impossible to determine whether phosphorylation of RyR2 itself was required for tetramer rearrangement and, if so, which residues were involved. To investigate this, and to further investigate our model, we used well-characterized homozygous transgenic mice, S2808A^+/+^ (Benkusky et al., 2007), S2814A^+/+^ (Chelu et al., 2009), S2814D^+/+^ (van Oort et al., 2010) and S2030A^+/+^ (Potenza et al., 2019) which have generated functional data to which our structural analyses could be compared. The specific residues were selected because their phosphorylation has been implicated in the β-adrenergic ‘fight or flight’ response, in elevated diastolic Ca^2+^ leak and in cardiac disease (Gaburjakova et al., 2020). S2808 and S2030 are phosphorylated by protein kinase A (PKA) and S2814 by calcium-calmodulin-dependent protein kinase II delta (CaMKIIδ) (Witcher et al., 1991; Marx et al., 2000; Wehrens et al., 2004; Xiao et al., 2005; Grimm et al., 2015; Potenza et al., 2019).

Mice were divided into groups whose hearts were perfused either with saline (control) or 300 nmol/L isoproterenol (Iso). Given that RyR2 cluster sizes may be quickly altered by phosphorylation (Asghari et al., 2020), the length of close apposition between the jSR and the t-tubule or surface membrane, the dyad, was measured from 2D transmission electron micrographs (Lavorato et al., 2015). The array’s organization was examined using our well-established dual-tilt electron tomography acquisition and analysis techniques (Asghari et al., 2014; Asghari et al., 2020).

The S2814D phosphomimetic mutant revealed a direct link between the state of the tetramer and the microarchitecture of the dyad, implying that phosphorylation of RyR2 itself initiates both an expansion of the dyad and a reorganization of the array, though the mechanisms are likely different. Iso produced the same structural changes in all of the WT mice and the S2808A and S2814A mutants, but not in the S2030A where the dyad did not expand and tetramer movement was crippled. All the mutants produced an abnormally large number of isolated tetramers indicating that each of the residues is important for maintaining inter-tetramer contact. The results showed that S2030 and S2808 are required for the normal β-adrenergic response, while S2814 is not. Using these same mouse models, published functional studies have supported these findings (van Oort et al., 2010; Respress et al., 2012; Ullrich et al., 2012; Potenza et al., 2019; Potenza et al., 2020).

## Materials and Methods

The experiments used ventricular myocytes from 10 to 16 week-old male C57BL/6J mice. Two hearts from each of RyR2-S2808A, RyR2-S2030A and their wild-type were the gift of Dr. H. Valdivia (Benkusky et al., 2007; Potenza et al., 2019), two hearts from each of RyR2-S2814A, RyR2-S2814D and their wild-type were the gift of Dr. X. Wehrens (Chelu et al., 2009). Hearts were extracted and hung on a Langendorff apparatus where they were perfused with either a physiological saline solution or with saline plus 300 mm/L Isoproterenol, for 2 min. They were then perfused with glutaraldehyde at a concentration of 4% in a 0.1 mmol/L cacodylate buffer, pH 7.4, for 5 minutes.

The left ventricular apex of each heart was cut into roughly 1mm X 1mm cubes using double edge razor blades. Post fixation, dehydration, infiltration and resin embedding were done as previously described (Asghari et al., 2009). Briefly, 4-5 cubes from each heart were post fixed in 2% OsO_4_ in 1mmol/L cacodylate buffer, then stained *en block* in 2% aquatic saturated uranyl acetate, after which it was gradually dehydrated in ethanol and embedded in a mixture of Spur and Epon. After 48h in a 60° C oven, samples were sorted and prepared for sectioning. Using a Leica UC7/FC7 microtome and a 35° diamond knife (DiATOME ultra, 2.5 mm), the embedded blocks were sectioned, about 80 nm thick for 2D imaging and 200 nm thick for 3D imaging. The sections were then placed on 2 mm x 1 mm Cu, Formvar coated slot grids (Electron Microscopy Sciences, Hatfield PA). After contrast staining, the samples were observed using a 200 kV Tecnai G2 TEM with a LaB6 filament (FEI, Hillsboro, OR). We examined two hearts for each of the mutant and WT mice; 2 blocks per heart, 4 sections per block and one section per grid. The same blocks were used for 2D and 3D EM. The total number of the dyads examined in each condition is indicated in Figure 2.

### Morphological Measurements

The Fiji program (NIH) was used for morphometric analysis of randomly collected 2D EM images from all phosphomutant and WT ventricular tissue. We measured the length of the dyad, defined as the region of close apposition between the jSR and surface or t-tubule membranes, in images at 30K magnification or higher (Lavorato et al., 2015). Supplementary Figure 1 displays details of how these measurements were performed. Samples were presented randomly to the individual measuring the lengths (S.S.) who was blinded as to their identity. The wild-type littermates were used as the controls; there was a single WT mouse for both the S2808A and S2030A mutants (WT 2808_2030), and the WT mice for the S2814A and WT S2814D mutants were pooled (WT 2814). All chemicals were purchased from Sigma-Aldrich (Oakville, ON) unless otherwise stated. Additional details of tissue preparation and EM acquisition are provided in Asghari *et al*. (Asghari et al., 2014; Asghari et al., 2020).

### Electron Tomography and Classifying RyR2 Positions

Each grid was examined on a dual-tilt electron microscope, a 200 kV Tecnai G2 (FEI, Hillsboro, OR), and relatively flat dyads were selected for imaging. We used Inspect3D to align the two tilt-series and Amira 6.4 (Thermo Fisher Scientific) to visualize the data sets, and employed no image processing steps other than a contrast stretch. Identification of the tetramers, their relative positions and measurement of the nearest neighbor distances were made using published methods (Asghari et al., 2014; Asghari et al., 2020).

Figure 1Ai displays a single plane from the tomogram acquired from a S2814A mouse heart. Each plane is outlined in a different colour; XY red, YZ green and XZ blue. Bisecting the dyad along the XY plane provided an *en face* view of the tetramers (Figure 1Aii). In most tomograms a single plane reveals limited information since the surface of the SR is uneven and not all the tetramers are either normal to the plane or visible. Individual XZ planes were therefore viewed individually through the dyad at 0.5 nm increments to best identify a RyR2 channel, then tilted in X and Y to identify its corners. That tetramer’s position and orientation were then mapped with a red box (27 x 27 nm) and the process was repeated for each tetramer. All the boxes were then displayed on a single plane from which the NND measurements were made (Figure 1Aiii). We select relatively flat junctions for tomographic analysis to enable a 2D projection of the 3D structure with limited distortion (Asghari et al., 2014).

Placement of the red boxes and calculating their centre-to-centre NND used a C++ program, ‘RyRFit’ written by one of the authors (DS). A detailed example of the process has been published (Asghari et al., 2020). Supplementary Figures 2 through 7 display images for the WT 2030_2808, WT 2814, S2808A, S2030A, S2814A and S2814D hearts, with and without 300 mmol/L Iso, respectively.

In Figure 1Aiv, the tetramers were colour-coded to indicate their relative positions. A tetramer was in a checkerboard arrangement (aqua) relative to its nearest neighbour if the sides were parallel, separated by less than 3 nm and overlapped by less than 18 nm 2/3 of its length. If those criteria were fulfilled but the overlap exceeded 18 nm, the tetramers were classified as side-by-side (orange). In those instances where the tetramers were farther apart than 3 nm, or their sides were not parallel, they were considered isolated (green) (Asghari et al., 2014). Figure 1B displays magnified views of tetramers in various configurations.

**Figure 1).**
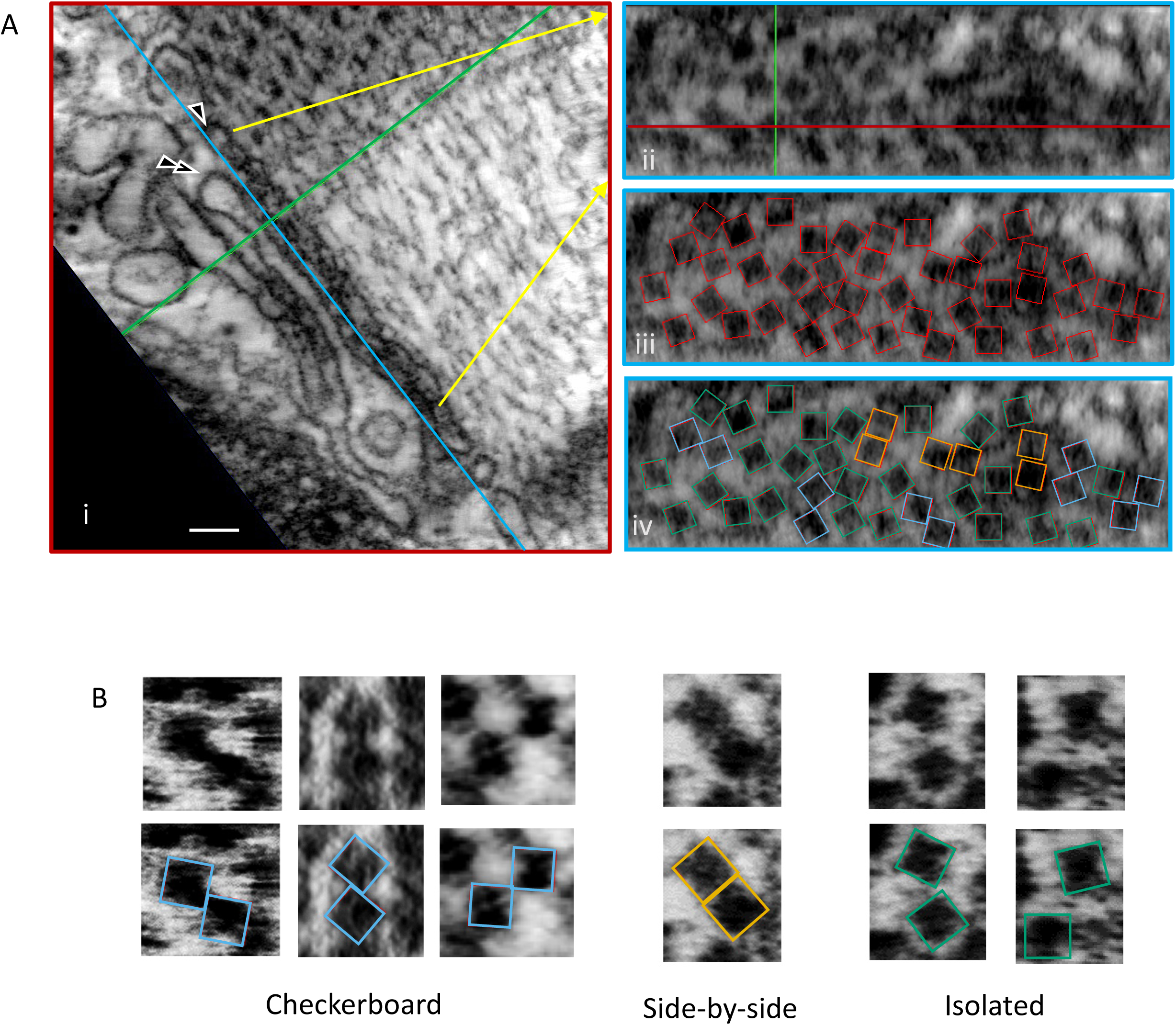
(A) A single image plane extracted from the dual-tilt tomogram of a mouse (S2814A) left ventricular myocyte fixed *in situ*. Different orthogonal planes from within the volume of the tomogram are indicated by different colors; XY in red, YZ in green and XZ in blue. The intersection point of all three planes lies within a single ryanodine receptor and the XZ plane (blue line) was positioned to parallel, as nearly as possible, the jSR (single arrowhead) and t-tubule (double arrowhead) membranes, but to be within the cleft and to bisect the ryanodine receptors. Scale bar is 100 nm (Ai). An *en face* view (XZ plane) of the indicated section (yellow arrows, Ai) of this junction (Aii). The position of the tetramers have been identified by a red box 27 x 27 nm (Aiii). Tetramers were colour-coded to indicate their orientation relative to their nearest-neighbour; checkerboard – aqua, side by side – orange, isolated – green. (B) Examples of the different tetramer-tetramer contacts.

### Statistical Analyses

Dyad lengths are presented as a violin plot. The data were analysed using hierarchical data analysis (HDA) to isolate the effects of genetic mutations and Iso from variations within a given experimental group, the cells and mice (Sikkel et al., 2017). This also removed the effects of pseudoreplication given that only two mice were available in each group. The data was not normally distributed, so similar to Sikkel we used the log transformation for analysis, followed by the Holm-Bonferroni correction(Holm, 1979). This is preferred to the Bonferroni correction which has a greater family-wise failure rate, is statistically weak and overly conservative (Olejnik et al., 1997). Tables 1 and 2, and Supplemental Table 1, have five columns; the first two show the comparison, the third, the probability calculated by Sikkel’s R program and the fourth and fifth, the results from the Holm-Bonferoni test. The p-values in column 3 were ranked from smallest to largest and a Holm-Bonferroni critical p-value, using α = 0.05, was calculated from the ranking; column 4. The comparisons showed a significant difference, column 5, if the actual p value was less than the critical p.

**Table 1.**
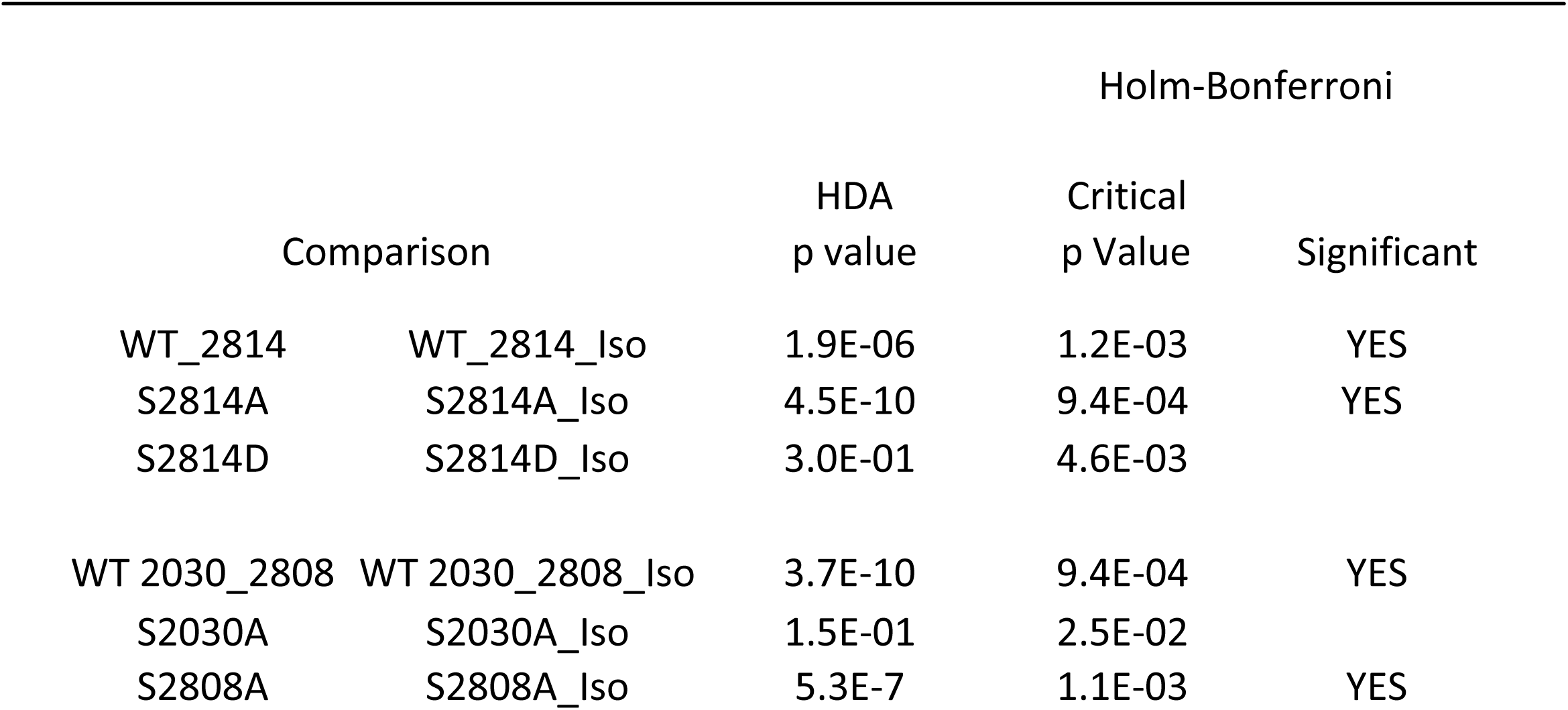
Effect of Isoproterenol on Dyad Length in Wild Type and Phosphomutant Mouse Ventricle

|  |  | Holm-Bonferroni |  |  |
| --- | --- | --- | --- | --- |
| Comparison |  | HDA<br>p value | Critical<br>p Value | Significant |
| WT_2814 | WT_2814_Iso | 1.9E-06 | 1.2E-03 | YES |
| S2814A | S2814A_Iso | 4.5E-10 | 9.4E-04 | YES |
| S2814D | S2814D_Iso | 3.0E-01 | 4.6E-03 |  |
| WT 2030_2808 | WT 2030_2808_Iso | 3.7E-10 | 9.4E-04 | YES |
| S2030A | S2030A_Iso | 1.5E-01 | 2.5E-02 |  |
| S2808A | S2808A_Iso | 5.3E-7 | 1.1E-03 | YES |

**Table 2.** S2814D: Pairwise Comparison of Dyad Lengths

|  |  | Holm-Bonferroni |  |  |
| --- | --- | --- | --- | --- |
| Comparison |  | HDA<br>p value | Critical<br>p Value | Significant |
| S2814D | WT 2814 | 9.4E-10 | 9.8E-04 | YES |
| S2814D | WT 2030_2808 | 6.2E-07 | 1.1E-03 | YES |
| S2814D | S2808A | 8.1E-05 | 1.4E-03 | YES |
| S2814D | S2814A | 4.7E-08 | 1.0E-03 | YES |
| S2814D | S2030A | 1.1E-10 | 9.1E-04 | YES |
| S2814D | WT_2814_Iso | 2.3E-01 | 2.2E-03 |  |
| S2814D | WT 2030_2808_Iso | 7.1E-01 | 5.0E-03 |  |
| S2814D | S2808A_Iso | 4.4E-01 | 5.6E-03 |  |
| S2814D | S2814A_Iso | 5.6E-01 | 3.3E-03 |  |
| S2814D | S2030A_Iso | 2.2E-08 | 1.0E-03 | YES |
| S2814D_Iso | WT 2030_2808_Iso | 1.3E-01 | 6.3E-03 |  |
| S2814D_Iso | WT_2814_Iso | 9.0E-01 | 2.2E-03 |  |
| S2814D_Iso | S2808A_Iso | 6.9E-02 | 7.1E-03 |  |
| S2814D_Iso | S2814A_Iso | 1.1E-01 | 3.6E-03 |  |
| S2814D_Iso | S2030A_Iso | 5.7E-05 | 1.4E-03 | YES |

The distribution of the tetramer centre-to-centre nearest-neighbour distances were compared using the non-parametric k-sample Anderson-Darling test (Scholz and Stephens, 1987) with values of α < 0.05 considered significant. All of the groups had significant differences within them. The Holm-Bonferroni correction was used for subsequent pair-wise comparisons within each group. We use the cumulative distribution function (CDF) when displaying distributions as this makes differences easier to visualize.

GraphPad Prism was used for the violin plot, histograms and CDFs using the Okabe-Ito colour palette to accommodate the colour blind (Katsnelson, 2021).

## Results

### Dyad Lengths

Superresolution immunofluorescence imaging indicated that RyR2 cluster sizes are significantly increased by a phosphorylation cocktail (Asghari et al., 2020), suggesting that physiological stimuli might also regulate the size of the dyads. To test that hypothesis, we measured the 2D length of mouse dyads in transmission electron micrographs. Figure 2A shows examples of dyadic clefts that had been perfused with saline or with saline plus 300 mmol/L isoproterenol (Iso); arrows show the end points of the measurements. The data is displayed in violin plots, Fig. 2B; the median, first and third quartiles are indicated with horizontal lines.

**Figure 2).**
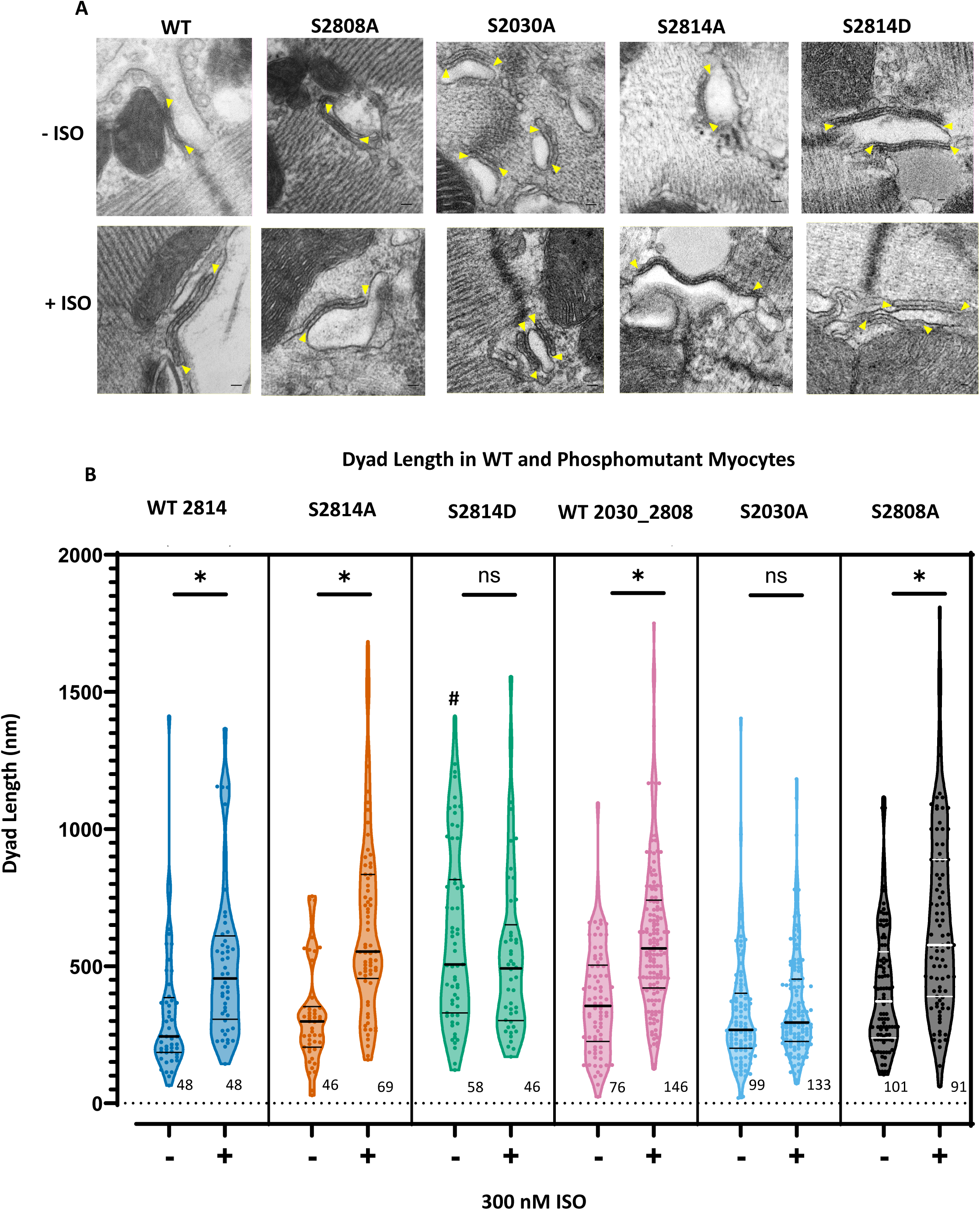
Dyad lengths. (A) TEM images under the indicated conditions, arrows indicate dyads’ extent (B) Violin plots of the dyad lengths. * indicates a significant difference between the given mutant and its ISO treated counterpart (p < 0.05). In the non-ISO treated hearts, dyads in the S2814D mouse were significantly longer than all others (#, p < 0.0013).

For clarity, the results of the hierarchical analysis are presented in their entirety in Supplemental Table 1. Selected comparisons are in Tables 1 and 2.

Table 1 shows the comparison between the dyad lengths of the saline treated mouse hearts and their Iso treated counterpart. Both WT mice showed highly significant increases in dyad length in response to Iso, as did the S2808A and S2814A myocytes. Notably, no changes were seen in either the S2814D or the S2030A mutants.

The effects of the S2814D mutation are highlighted in Table 2. At rest, its dyad was significantly longer than the others, and with the exception of S2030A it was no different from those that had been treated with Iso. Perfusing the S2814D hearts with Iso had no further effect on the length of their dyads.

Measuring the length of the dyad, following the curve of the membrane, is a simple measurement, but it produces clear results. The dyad cross-sections are a 2-D view of a 3-D volume that extends in both directions in the Z dimension. We don’t know where our cross sections lie relative to the dyad’s maximum diameter, nor how they are oriented with respect to it. Thus, the measured cross-sections represent cuts at random positions and angles through the junction. For these cuts to produce a significant change in length, it is highly likely there was an expansion of the dyad’s area, not just its length.

### Tetramer Orientations and Nearest-Neighbour Distances

#### Wild Types

Figures 3Ai and 3Bi display the RyR2 NND and orientation histograms obtained from the two WT mice; images of the tomograms and tetramer placements for saline and Iso perfused hearts are in Supplementary Figs. 1 and 2. The histograms were bimodal, with one mode (∼28 nm) dominated by tetramers in the side-by-side (orange) orientation and the second (∼34 nm) by those in a checkerboard (aqua). A few tetramers were classified as isolated (green). The percentage of tetramers in each orientation and the number of tomograms and tetramers examined are listed in Table 3.

**Table 3.** Tetramer Orientations (%)

| Mouse | Checkerboard | Side-by-Side | Isolated | Tomograms | Tetramers | Mice |
| --- | --- | --- | --- | --- | --- | --- |
| WT S2030-S2808 | 46.7 | 38.3 | 15.0 | 3 | 60 | 2 |
| WT S2030-S2808 + Iso | 81.3 | 12.0 | 6.7 | 3 | 75 | 2 |
| S2808A | 39.3 | 44.0 | 16.7 | 4 | 84 | 2 |
| S2808A + Iso | 63.8 | 12.3 | 23.9 | 7 | 138 | 2 |
| S2030A | 35.0 | 32.0 | 33.0 | 6 | 103 | 2 |
| S2030A + Iso | 44.7 | 29.8 | 25.4 | 6 | 114 | 2 |
| WT S2814 | 53.9 | 33.1 | 13.0 | 17 | 369 | 2 |
| WT S2814 + Iso | 76.3 | 7.9 | 15.8 | 13 | 227 | 2 |
| S2814A | 45.0 | 17.4 | 37.6 | 4 | 109 | 2 |
| S2814A + Iso | 69.8 | 14.4 | 15.8 | 9 | 139 | 2 |
| S2814D | 60.7 | 11.9 | 27.4 | 6 | 135 | 2 |
| S2814D + Iso | 78.6 | 7.1 | 14.3 | 6 | 84 | 2 |

Perfusing the WT mouse hearts with Iso for two minutes resulted in unimodal histograms with peaks at ∼34 nm (Figs 3Aii and Bii). These changes were accompanied by a large increase in the proportion of tetramers in the checkerboard orientation and a correspondingly large decrease in those that were side by side, while those in the isolated configuration remained few in number (Table 3).

Both WT responded to Iso with an increase in their tetramer NNDs which is reflected in a significant rightward shift of their CDFs. Neither the saline perfused WTs nor the Iso treated were significantly different from each other, Fig. 3C, Table 4. These histograms and tetramer orientations were similar to those recorded from rat and human(Asghari et al., 2014; Asghari et al., 2020), resulting in comparable CDFs, Fig. 3D.

**Figure 3).**
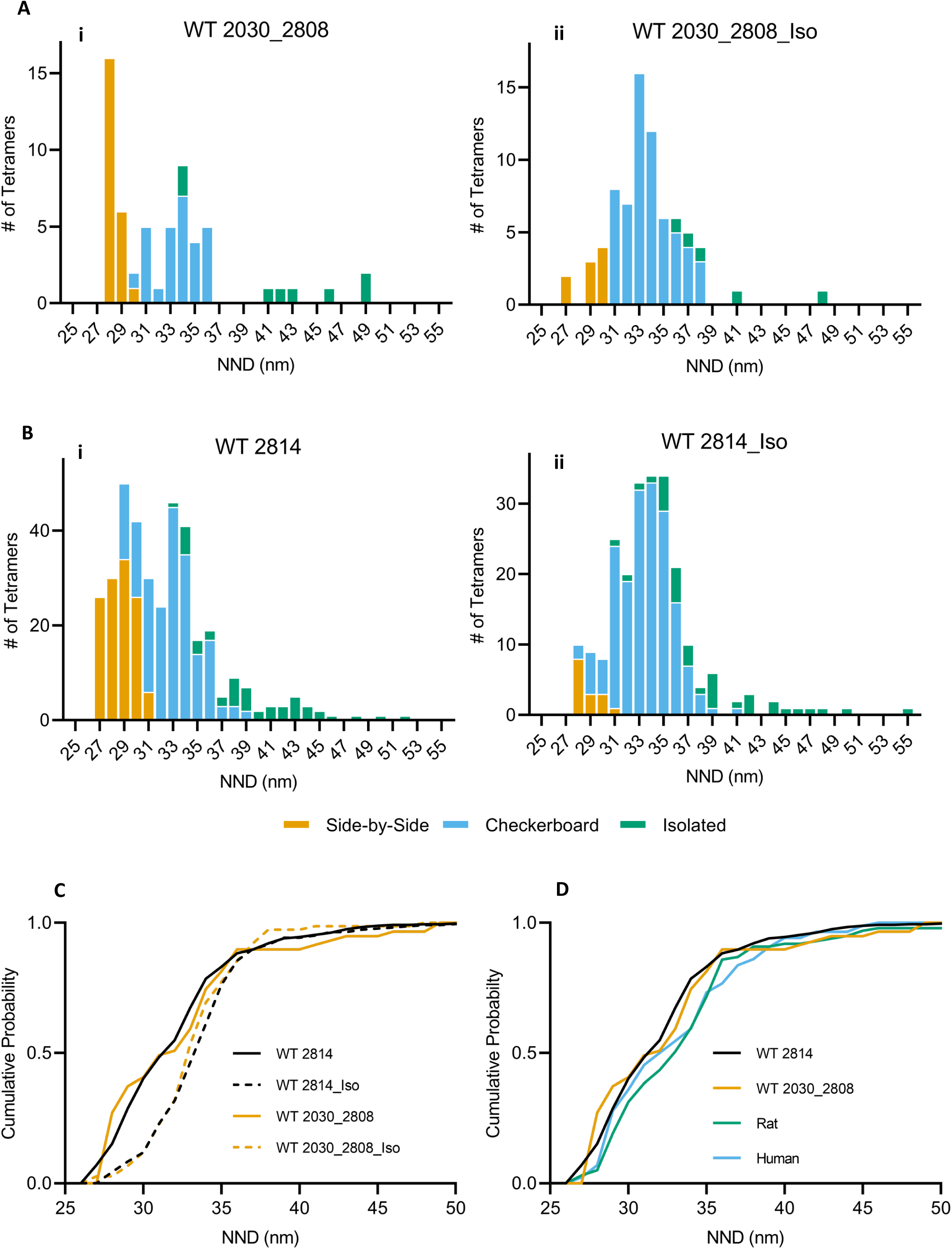
Nearest-neighbour distances of RyR2 acquired from WT S2030_S2808 (A) and S2814 (B) mice, at rest (i) and from mice whose hearts had been treated with ISO (ii). C) CDF of tetramer NND acquired from WT mice hearts at rest and from those treated with ISO D) CDF of tetramer NND comparing WT mice with rat and human.

**Table 4.** CDF Comparisons

| Group | Paired Comparison |  | p | Critical<br>p value | Significant |
| --- | --- | --- | --- | --- | --- |
| WT Mice | WT 2814 | WT 2814_Iso | 1.5E-12 | 8.0E-03 | YES |
|  | WT 2814 | WT 2030_2808_Iso | 2.8E-05 | 1.0E-02 | YES |
|  | WT 2814_Iso | WT 2030_2808 | 8.1E-05 | 1.3E-02 | YES |
|  | WT 2030_2808 | WT 2030_2808_Iso | 1.7E-03 | 1.7E-02 | YES |
|  | WT 2814 | WT 2030_2808 | 2.1E-01 | 2.5E-02 |  |
|  | WT 2814_Iso | WT 2030_2808_Iso | 5.5E-01 | 5.0E-02 |  |
| Species | WT 2814 | Rat | 3.4E-03 | 8.0E-03 | YES |
|  | WT 2814 | Human | 1.3E-02 | 1.0E-02 |  |
|  | WT 2030_2808 | Rat | 3.6E-02 | 1.3E-02 |  |
|  | WT 2030_2808 | Human | 4.4E-02 | 1.7E-02 |  |
|  | WT 2814 | WT 2030_2808 | 2.1E-01 | 2.5E-02 |  |
|  | Rat | Human | 5.7E-01 | 5.0E-02 |  |
| S2808A | WT 2030_2808 | S2808A_Iso | 1.6E-03 | 8.0E-03 | YES |
|  | S2808A | S2808A_Iso | 1.7E-03 | 1.0E-02 | YES |
|  | WT 2030_2808 | WT 2030_2808_Iso | 1.7E-03 | 1.3E-02 | YES |
|  | WT 2030_2808_Iso | S2808A | 2.2E-03 | 1.7E-02 | YES |
|  | WT 2030_2808_Iso | S2808A_Iso | 4.6E-02 | 2.5E-02 |  |
|  | WT 2030_2808 | S2808A | 2.2E-01 | 5.0E-02 |  |
| S2030A | WT 2030_2808 | WT 2030_2808_Iso | 1.7E-03 | 8.0E-03 | YES |
|  | WT 2030_2808 | S2030A_Iso | 2.3E-03 | 1.0E-02 | YES |
|  | S2030A | S2030A_Iso | 2.9E-03 | 1.3E-02 | YES |
|  | WT 2030_2808_Iso | S2030A | 5.7E-03 | 1.7E-02 | YES |
|  | WT 2030_2808_Iso | S2030A_Iso | 3.1E-02 | 2.5E-02 |  |
|  | WT 2030_2808 | S2030A | 5.3E-01 | 5.0E-02 |  |
| S2814A | WT 2814 | S2814A | 4.8E-13 | 8.0E-03 | YES |
|  | WT 2814 | WT 2814_Iso | 1.5E-12 | 1.0E-02 | YES |
|  | WT 2814 | S2814A_Iso | 7.0E-09 | 1.3E-02 | YES |
|  | WT 2814_Iso | S2814A | 4.5E-03 | 1.7E-02 | YES |
|  | S2814A | S2814A_Iso | 2.7E-02 | 2.5E-02 |  |
|  | WT 2814_Iso | S2814A_Iso | 2.9E-01 | 5.0E-02 |  |
| S2814D | WT 2814 | WT 2814_Iso | 1.5E-12 | 8.0E-03 | YES |
|  | WT 2814 | S2814D | 1.2E-04 | 1.0E-02 | YES |
|  | WT 2814 | S2814D_Iso | 1.7E-04 | 1.3E-02 | YES |
|  | WT 2814_Iso | S2814D | 3.3E-02 | 1.7E-02 |  |
|  | WT 2814_Iso | S2814D_Iso | 8.8E-02 | 2.5E-02 |  |
|  | S2814D | S2814D_Iso | 7.0E-01 | 5.0E-02 |  |

#### Phosphomutants

Figures 4A (S2808A), 4B (S2030A), 5A (S2814A) and 5B (S2814D), display NND histograms from myocytes whose hearts had been perfused with saline (i) or saline plus Iso (ii), and the corresponding CDFs (iii). The tetramer orientations, and the number of tetramers and tomograms examined, are in Table 3. Images of the tomograms and tetramer placements for the saline and Iso perfused hearts are in Supplementary Figs. 3 (S2808A), 4 (S2030A), 5 (S2814A) and 6 (S2814D).

**Figure 4).**
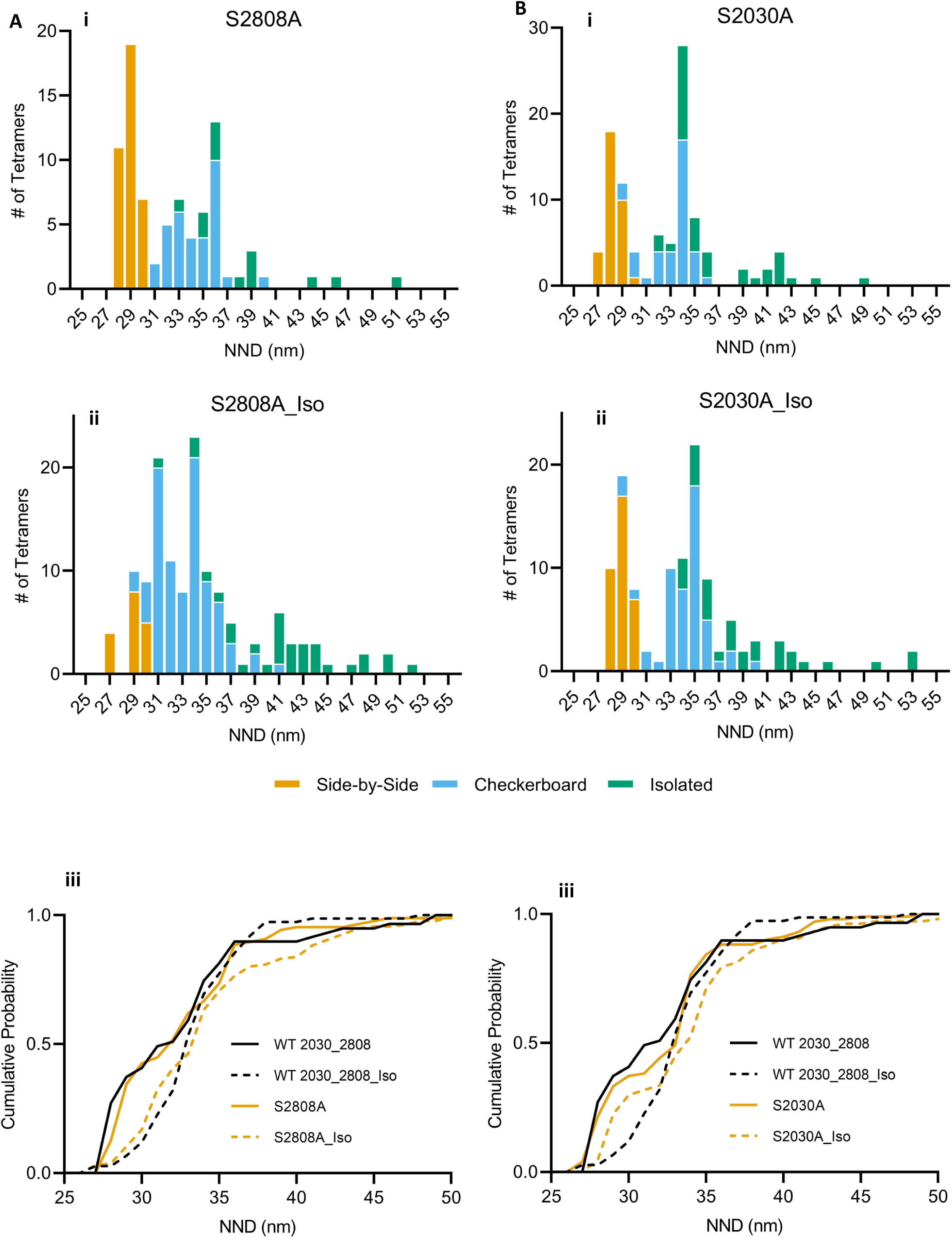
Nearest-neighbour distances of RyR2 acquired from S2808A (A) and S2030A (B) mice, at rest (i) and from mice whose hearts had been treated with ISO (ii). C) CDF of tetramer NND acquired from WT 2030_2808 and S2808A hearts at rest and from those treated with ISO. D) Same as C for S2030A.

##### S2808A

The NND distribution (Fig. 4Ai), tetramer orientations (Table 3) and the CDF (Fig. 4Aiii, Table 4) were identical to that of the WT.

On application of Iso, the CDFs were not significantly different, but the orientations showed a significant difference in their response. Treatment with Iso caused a large reduction in the proportion of isolated tetramers in the WT, from 15.0% to 6.7%. Iso had the opposite effect in this mutant, with the proportion of isolated tetramers increasing from 16.7% to 23.9%, with an associated drop in the checkerboard from 81.3% to 63.8%.

##### S2030A

This NND distribution was bimodal (Fig 4Bi), comparable to WT, and the CDFs were not significantly different (Table 4). Despite this the tetramer orientations were very different, with double the proportion of isolated, and roughly an equal distribution between checkerboard, side-by-side and isolated (Table 3).

The Iso response showed a significant change in the CDF (Table 4) compared to the WT, although the tetramers with a NND < 32 nm seemed to have a muted response, probably because the proportion of side-by-side tetramers was largely unchanged. There was only a small increase in the checkerboard and a comparable decrease in the isolated.

##### S2814A

Although the NND distribution of this mutant was bimodal (Figure 5Ai), the modes corresponded to unusual orientations, with one at 31 nm composed largely of checkerboard and the second at 35 nm with roughly equal proportions of checkerboard and isolated. The number of isolated tetramers was three times that of the WT (37.6% vs. 13.0%), the side-by-side was half (17.4% vs 33.1%), and there were fewer checkerboard (45.0% vs 53.9%). The result of these changes was a highly disorganized array (Fig. 1A).

The NND distribution of the Iso treated myocytes appeared not as broad and was unimodal with a peak between 32 and 34 nm, smaller than the WT (Fig. 5Aii). This change reflected a 60% drop in the isolated tetramers, to 14.4%, particularly those at NND > 40 nm, and a corresponding rise in the checkerboard (69.8%), while the side-by-side were unchanged (Table 3).

The rightward shift of the CDF in response to Iso seen in WT mice and rats reflects a movement of side-by-side tetramers to a largely checkerboard configuration. In this case, the proportion of side-by-side tetramers in the S2814A mice was low, and changed minimally in response to ISO, so that part of the CDF was already shifted and there was no further change. The result was that the S2814A CDF was significantly different from the WT 2814, but not from the S2814A_Iso (Table 4).

##### S2814D

The NND distribution had a single peak at ∼33 nm reflecting the small proportion of tetramers in the side-by-side orientation, with a large increase in the number of isolated and a small increase of the checkerboard (Fig 5Bi Table 3). The sharp reduction in the side-by-side is similar to Iso perfused WT hearts, and in rats after exposure to a phosphorylation cocktail (Asghari et al., 2014; Asghari et al., 2020), and is consistent with S2814D being a phosphomimetic. As a result, the CDF was shifted to the right and was not significantly different from the WT treated with Iso (Table 4).

Iso produced little change in the NND distribution, Fig 5Bii, resulting in no significant difference between the CDFs of S2814D and S2814D_Iso or between S2814D_Iso and WT 2814_Iso. There was a very large increase in the number of tetramers in the checkerboard orientation, a large decrease in the isolated and a minimal change in the side-by-side (Table 3).

## Discussion

### Dyad Lengths

We have found that the size of the dyad in WT and mutant mouse ventricular myocytes, with the notable exceptions of S2814D and S2030A, was significantly increased by a two-minute exposure to 300 mmol/L Iso.

The lengths of the dyads in the S2814D mutant were equivalent to those of WT + Iso for the saline perfused hearts. That the expansion of the dyad is produced by a single mutation in RyR2 implies a direct link between the state of the tetramer and the architecture of the membrane in which it is embedded. The lack of further response of the S2814D + Iso cells suggests that the effect of phosphorylation of other sites on RyR2 is not additive and that the expansion derived from S2814D alone is near maximal.

In considering whether the changes in dyad length that we attribute to the mutations alone could be due to changes in the expression of RyR2 or its associated proteins, we note that within the limits of detection of the Western blotting technique, there were no changes in the expression of RyR2, Cav1.2 or the Na^+^/Ca^2+^ exchanger in the S2814D mice (van Oort et al., 2010). If there had been undetectable changes in expression this would not explain the lack of response to Iso. The expansion of the dyads of WTs, S2080A and S2814A in response to Iso is too rapid to be explained by changes in protein expression. This data taken together with the S2814D behaviour, suggests a connection between the RyR2 and the jSR which quickly affects the size of the dyad.

We do not know what the link might be and whether it is direct or via RyR2’s attendant proteins, although there is evidence that the phosphomimetic S2814D has a changed RyR2 proteome (Chiang et al, 2021). Notably, the application of Iso produced no expansion of the dyad in the S2030A mouse. Whether the lack of response is due to interrupting this link or through another mechanism is unknown.

Direct, real time, visualization of the SR and t-tubule in living murine myocytes has demonstrated that dyads are highly mobile structures that are continuously and rapidly changing in number, shape, size and position (Vega et al., 2011; Drum et al., 2020). Within minutes, dyads were seen to form, disappear and to split into smaller pieces; a movement driven in part by the microtubule motor proteins kinesin-1 and dynein, both of which are subject to regulation by a host of post translational modifications that alter the direction of movement, the transit time and the binding and unbinding of their cargo (Barlan and Gelfand, 2017; Kumari, 2022). In this paradigm, the size of the dyads we observe in our fixed tissues are snapshots of this dynamic process. Iso, which activates PKA and, through Epac 2, CamKII (Pereira et al., 2013), could shift the equilibrium to one in which larger more stable dyads are favoured. Recent research has also demonstrated that motor-cargo adaptors, including integral membrane proteins, can recruit kinesin and dynein to an organelle to regulate its transport (Cross and Dodding, 2019). Our results with the S2814D and S2030A mice suggest that the phosphorylation status of RyR2 is a central element in these processes either directly, or indirectly by altering its binding partners (Chiang et al., 2021).

The nanoscopic domain formed by the dyad is a critical determinant of excitation-contraction coupling. Within its femtoliter volume, diffusion is restricted to enable the ion channels, signaling molecules and regulatory proteins on the apposing membranes to control SR Ca^2+^ release independently of reactions within the bulk myoplasm. Changing its size, as we have observed, would be expected to change the number and spatial distribution of the molecules therein which would impact Ca^2+^ dynamics, the probability of Ca^2+^ spark formation (Iaparov et al., 2021) and the efficacy of excitation-contraction coupling.

#### Organization of the Tetramer Array

##### WT Mice

Under basal conditions, the tetramers in the WT mouse ventricular dyads were distributed similarly to each other and to rats and humans, with a bimodal CDF of their NND reflecting a mixture of checkerboard and side-by-side orientations. Most of the tetramers were in contact with one or more neighbours, though a few were isolated (Figure 3D).

Cabra *et al* (Cabra et al., 2016) have observed tetramer-tetramer interactions using cryo-electron microscopy to examine purified RyR2 laid on a carbon coated grid. Despite the near infinite number of possible interactions between tetramers under these experimental conditions, they saw only a few, distinct, configurations termed adjoining and oblique, which correspond well to our side-by-side and checkerboard respectively. The proportion in each configuration could be adjusted by changing the Ca^2+^ concentration, indicating that the interactions may be sensitive to physiologically relevant parameters.

That similar tetramer-tetramer contacts have been visualized *in vivo* in three different species and *in vitro* using vastly different techniques, suggests that inter-tetramer interaction is real and an integral part of the array’s normal structure and function. That conclusion is further supported by our observations using dSTORM immunofluorescence microscopy, which also demonstrates tetramer-tetramer contacts similar to those we have described (Scriven et al., 2023).

Treatment with Iso resulted in fewer side-by-side tetramers and an increase in the checkerboard (Figures 2Aii, Bii and Table 3), which was reflected by a rightward shift of the CDF (Figure 2C). We observed the same changes in tetramer orientation and NND CDF from rat myocytes that were enzymatically dissociated, permeabilized and treated with a phosphorylation cocktail that activated numerous kinases and inhibited the major phosphatases (Asghari et al., 2014; Asghari et al., 2020). Taken together, these results show that activation of the β-adrenergic receptor induces tetramer reorganization as well as expansion of the dyad, indicating that both are part of the normal fight-or-flight response.

In all of our experiments, changes in the CDF and tetramer orientation were accompanied by a change in the length of the dyad, with the exception of S2030A. In this mutant, some of the side-by-side and isolated tetramers reoriented into a checkerboard configuration resulting in a significant shift of the CDF, but the length of the dyad was unchanged. This implies that changes in the size of the dyad and the tetramers’ relative positions and orientations are the result of different processes.

##### Phosphomutant Mice

All of the substitutions produced significant changes in the organization of the array compared to their WT counterparts. In particular, they had an abnormally large number of isolated tetramers either after treatment with Iso (S2808A), before Iso (S2814A, S2814D) or both before and after (S2030A), indicating that all of these residues are involved in the normal structure of the array and in its restructuring in response to β−adrenergic stimulation (Table 3). How the mutations affect the tetramer-tetramer contacts or the long-range allosteric interactions required to open the channel (Peng et al., 2016) is unknown.

The majority of the RyR2 in the S2814D mutant were in a checkerboard configuration, which combined with the low number of side-by-side, and the rightward shift in the CDF, was close to the expected configuration of the array after treatment with Iso (Figure 5B, Table 3). This suggests that phosphorylation of the tetramer itself affects both the size of the dyad and the organization of the array, but does not preclude these events being mediated by different processes as suggested by the results from the 2030A mouse. Treating this mouse with Iso produced a further shift in the tetramers’ organization, becoming indistinguishable from WT + Iso indicating that phosphorylation of other residues, or downstream targets, is required to complete the transition.

**Figure 5).**
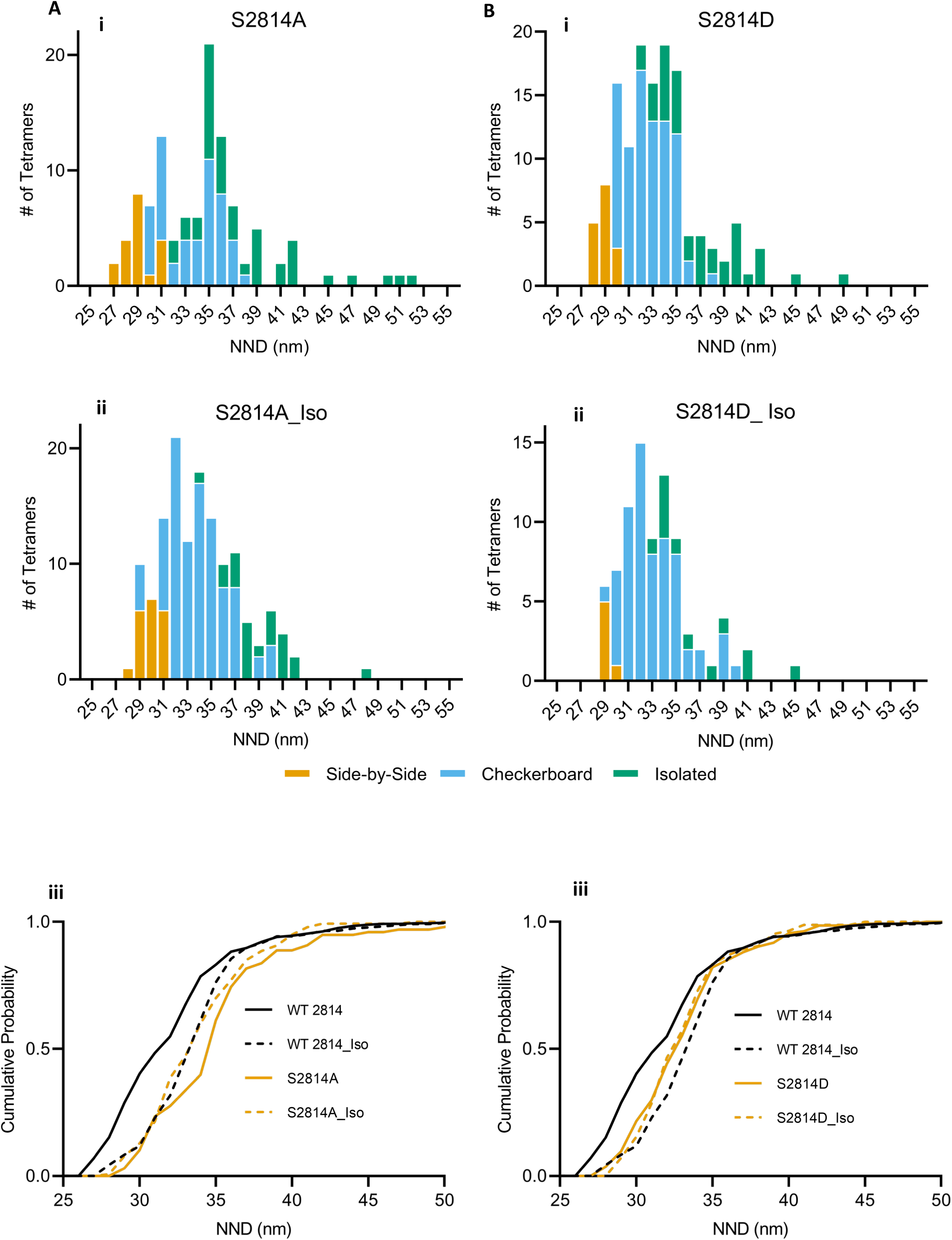
Nearest-neighbour distances of RyR2 acquired from S2814A (A) and S2814D (B) mice, at rest (i) and from mice whose hearts had been treated with ISO (ii). C) CDF of tetramer NND acquired from WT 2814 and S2814A hearts at rest and from those treated with ISO. D) Same as C for S2814D.

The position of the CDF in the S2814A mouse was comparable to WT + Iso and typical of an array with a much higher proportion of tetramers in the checkerboard configuration (Figure 5A, Table 3). In this case, it was due to a large number of isolated tetramers that were located at a sufficient distance from each other to shift the CDF to a significantly longer NND. The result was a highly disordered array (Figure 1). Like S2814D, treating this mutant with Iso resulted in an array that closely resembled WT + Iso.

We conclude that although phosphorylation of S2814 alone mimics β−adrenergic stimulation, the S2814A results suggest this residue is not necessary to elicit a response. The limited movement of the CDF and change in tetramer reorientation in response to Iso in the S2030A mutant, and the abnormal proportion of isolated tetramers following Iso in the S2808A, suggest that both of these residues are needed.

## Structure and Function

The conclusions derived from the structural analyses, that S2808 and S2030 are necessary for a normal β-adrenergic response, while S2814 is not, concurs with results from functional studies that have used the same transgenic mouse models. Briefly, these analyses have demonstrated that with matched Ca^2+^ current, Ca^2+^ transient and SR Ca^2+^ load, Iso produces less SR Ca^2+^ release from S2030A myocytes than from WT, resulting in a significantly lower ECC gain (Potenza et al., 2019). In the S2808A mutant, Iso results in a loss of intraluminal Ca^2+^ sensitivity and spatiotemporal synchronization of RyR2 release relative to WT(Ullrich et al., 2012; Potenza et al., 2020) but without affecting ECC gain (Benkusky et al., 2007; Ullrich et al., 2012). Both of these residues are therefore necessary for a normal β-adrenergic response. Conversely, S2814A mice appear to have normal ECC parameters in response to Iso (Grimm et al., 2015; Baier et al., 2021), so phosphorylation of this residue is not required.

Our structural analysis of the S2814D dyads suggested that the pseudophosphorylation was a gain-of-function mutation, a condition that has been observed (van Oort et al., 2010). They reported that this mutant has a high Ca^2+^ spark rate, despite a significantly reduced SR Ca^2+^ load. This change is pro-arrhythmogenic and has been linked to elevated diastolic Ca^2+^ leak in both patients and animals in heart failure (Ai et al., 2005; Ling et al., 2009) suggesting prolonged phosphorylation of this residue is pathological (Grimm et al., 2015).

The presumptive mechanism linking RyR2 phosphorylation to cardiac function are changes in single channel gating. Such effects have been recorded from isolated channels in artificial bilayers, but the magnitude of the changes in open probability do not match those seen *in vivo* (Benkusky et al., 2007; Xiao et al., 2007; van Oort et al., 2010; Potenza et al., 2019). This difference may be explained, in part, by the direct link we have identified between the state of the tetramer and the jSR, which is absent with purified RyR2 and unlikely to be preserved by heavy SR vesicles painted onto an artificial bilayer. Tetramer-tetramer contacts, also routine *in vivo*, are seldom observed *in vitro* and are important as our conceptual model postulates that inter-tetramer contacts modify channel gating, which has been observed with both RyR2 (Marx et al., 2001) and RyR1 (Porta et al., 2012).

We have compared the changes in the Ca^2+^ spark rate of the mutants relative to WT, both with and without Iso (or CaMKII in the case of S2814D), to our model’s predictions (Table 5). All of our predictions are in agreement with experimental observations with the exception of S2814A. Given the low proportion of side-by-side tetramers both at rest and after Iso we predicted a high Ca^2+^ spark rate under both conditions, which was not observed. At rest, the spark rate is comparable to WT (Uchinoumi et al., 2016) but after Iso the SR Ca^2+^ leak rate/Ca^2+^ load (a combination of Ca^2+^ spark and non-spark diastolic release) while significantly elevated in WT, is unchanged or even slightly lower in this mutant (Baier et al., 2021).

**Table 5.** Change in CaSpF in phosphomutant myocyte compared to WT: Model prediction vs experimental observation

|  | Control |  |  | Phosphorylated |  |
| --- | --- | --- | --- | --- | --- |
|  | Model | Observed |  | Model | Observed |
| S2030A(Potenza et al., 2019) | ↓ | ↓ |  | ↓ | ↓ |
| S2808A(Potenza et al., 2020) | ↔ | ↔ |  | ↑ | ↑ |
| S2814D (van Oort et al., 2010; Uchinoumi et al., 2016) | ↑ | ↑ |  | ↔ | ↔ |
| S2814A(Baier et al., 2021) | ↑ | ↔ |  | ↑ | ↔ |

The relative change in $\text{Ca}^{2+}$ spark frequency of the indicated phosphomutant myocyte relative to its WT is indicated by an arrow. The columns compare the change predicted by the model to the observed result for myocytes at rest (Control) and after phosphorylation by treatment with either Iso (S2030A, S2808A and S2814A) or CaMKII (2814D).

Despite the fact that our original model did not take into account the isolated tetramers, which are present in large numbers in the mutants, our model is successful in predicting the behaviour of WT rats and mice, and the majority of the mutants (Table 5). The sole exception, S2814A, cannot be explained with our model unless we assume that this mutation interrupts positive allosteric interaction associated with the checkerboard configuration in some way. This could prevent the increase in spark frequency predicted by the model and align it with the experimental data, but lacks evidence and remains speculation.

## Conclusions

We have found that both the size of the dyad and the arrangement of the tetramers are dynamically altered by application of the β-adrenergic receptor agonist, isoproterenol. The changes induced by the phosphomutant S2814D implied a link between the state of the tetramer, the size of the dyad and the tetramer organization.

In agreement with functional data, we found that S2808 and S2030 are necessary for the β-adrenergic response, while S2814 may not be. The disposition of the tetramers, and the correlation of structure with function both at rest and in response to Iso, suggests that tetramer-tetramer contacts plays an important functional role. We have proposed that the contacts modify the tetramers’ gating parameters, possibly by changing allosteric interactions.

In this paper and its companion (Scriven et al., 2023) we have observed comparable tetramer distributions, both on the surface and in the interior, despite different methodologies and fixation protocols. Both indicate the tetramers are in an irregular, but non-random, distribution with numerous tetramer-tetramer contacts. We found, separately, that isoproterenol produces significant and rapid changes in the structure. While our results are snapshots of the state of the dyad before and after the application of an agonist, the implication is that the structure is continuously shifting in response to changes in the local environment.

## Acknowledgements

E.D.W.M. acknowledges a grant from the Canadian Institutes of Health Research (148527). H.H.V. is funded by a grant from the US National Institutes of Health (R01-HL055438). X.H.T.W. is funded by grants from the US National Institutes of Health (HL089698, HL147108, HL153350). The authors declare no competing financial interests.

## Author Contributions

**Parisa Asghari**: Conceptualization, Resources, Investigation, Formal Analysis, Data Curation, Visualization, Writing – Original Draft, Writing – Review & Editing, **David R.L. Scriven**: Software, Formal Analysis, Visualization, Writing – Original Draft, Writing – Review & Editing. **Saba Shahrasebi**: Investigation. **Hector H. Valdivia**: Resources, Writing – Review & Editing. **Katherina M, Alsina**: Resources. **Xander H.T. Wehrens**: Resources, Writing – Review & Editing. **Edwin D.W. Moore**: Conceptualization, Formal Analysis, Visualization, Writing – Original Draft, Writing – Review & Editing, Funding Acquisition, Supervision.

## Abbreviations

CDF: Cumulative Distribution Function
ISO: Isoproterenol
NND: Nearest neighbor distance
RyR2: Type II Ryanodine Receptors

