## Supplemental_Figures for "PHOSPHORYLATION OF RyR2 SIMULTANEOUSLY EXPANDS THE DYAD AND REARRANGES THE TETRAMERS"

#### Supplemental Table 1

| Comparison | | HDA p value | Holm-Bonferroni Critical p value | Holm-Bonferroni Null Hypothesis |
|---|---|---|---|---|
| WT_2814 | WT_2814_Iso | 1.855E-06 | 0.00122 | REJECT |
| WT_2814 | S2814A | 6.079E-01 | 0.00179 | |
| WT_2814 | S2814A_Iso | 3.518E-12 | 0.00083 | REJECT |
| WT_2814 | S2814D | 9.363E-10 | 0.00098 | REJECT |
| WT_2814 | S2814D_Iso | 1.855E-06 | 0.00125 | REJECT |
| WT_2814 | WT 2030_2808 | 9.459E-02 | 0.00185 | |
| WT_2814 | WT 2030_2808_Iso | 1.265E-13 | 0.00079 | REJECT |
| WT_2814 | S2030A | 6.630E-01 | 0.00192 | |
| WT_2814 | S2030A_Iso | 1.005E-01 | 0.00200 | |
| WT_2814 | S2808A | 4.351E-03 | 0.00156 | |
| WT_2814 | S2808A_Iso | 4.133E-13 | 0.00081 | REJECT |
| WT_2814_Iso | S2814A | 3.038E-05 | 0.00132 | REJECT |
| WT_2814_Iso | S2814A_Iso | 6.947E-02 | 0.00208 | |
| WT_2814_Iso | S2814D | 2.288E-01 | 0.00217 | |
| WT_2814_Iso | S2814D_Iso | 8.960E-01 | 0.00227 | |
| WT_2814_Iso | WT 2030_2808 | 3.859E-04 | 0.00152 | REJECT |
| WT_2814_Iso | WT 2030_2808_Iso | 8.006E-02 | 0.00238 | |
| WT_2814_Iso | S2030A | 7.526E-07 | 0.00114 | REJECT |
| WT_2814_Iso | S2030A_Iso | 5.782E-05 | 0.00139 | REJECT |
| WT_2814_Iso | S2808A | 1.194E-02 | 0.00172 | |
| WT_2814_Iso | S2808A_Iso | 4.192E-02 | 0.00250 | |
| S2814A | S2814A_Iso | 4.475E-10 | 0.00094 | REJECT |
| S2814A | S2814D | 4.707E-08 | 0.00104 | REJECT |
| S2814A | S2814D_Iso | 2.802E-05 | 0.00128 | REJECT |
| S2814A | WT 2030_2808 | 2.815E-01 | 0.00263 | |
| S2814A | WT 2030_2808_Iso | 5.080E-11 | 0.00088 | REJECT |
| S2814A | S2030A | 8.754E-01 | 0.00278 | |
| S2814A | S2030A_Iso | 3.202E-01 | 0.00294 | |
| S2814A | S2808A | 2.653E-02 | 0.00313 | |
| S2814A | S2808A_Iso | 8.411E-11 | 0.00089 | REJECT |
| S2814A_Iso | S2814D | 5.573E-01 | 0.00333 | |
| S2814A_Iso | S2814D_Iso | 1.060E-01 | 0.00357 | |
| S2814A_Iso | WT 2030_2808 | 4.505E-09 | 0.00100 | REJECT |
| S2814A_Iso | WT 2030_2808_Iso | 7.563E-01 | 0.00385 | |
| S2814A_Iso | S2030A | 7.936E-14 | 0.00078 | REJECT |

| | | | | |
|---|---|---|---|---|
| S2814A_Iso | S2030A_Iso | 3.336E-11 | 0.00086 | REJECT |
| S2814A_Iso | S2808A | 1.813E-06 | 0.00119 | REJECT |
| S2814A_Iso | S2808A_Iso | 8.598E-01 | 0.00417 | |
| S2814D | S2814D_Iso | 3.018E-01 | 0.00455 | |
| S2814D | WT 2030_2808 | 6.198E-07 | 0.00111 | REJECT |
| S2814D | WT 2030_2808_Iso | 7.090E-01 | 0.00500 | |
| S2814D | S2030A | 1.101E-10 | 0.00091 | REJECT |
| S2814D | S2030A_Iso | 2.204E-08 | 0.00102 | REJECT |
| S2814D | S2808A | 8.100E-05 | 0.00143 | REJECT |
| S2814D | S2808A_Iso | 4.376E-01 | 0.00556 | |
| S2814D_Iso | WT 2030_2808 | 3.473E-04 | 0.00147 | REJECT |
| S2814D_Iso | WT 2030_2808_Iso | 1.269E-01 | 0.00625 | |
| S2814D_Iso | S2030A | 8.417E-07 | 0.00116 | REJECT |
| S2814D_Iso | S2030A_Iso | 5.705E-05 | 0.00135 | REJECT |
| S2814D_Iso | S2808A | 1.005E-02 | 0.00167 | |
| S2814D_Iso | S2808A_Iso | 6.870E-02 | 0.00714 | |
| WT 2030_2808 | WT 2030_2808_Iso | 3.706E-10 | 0.00093 | REJECT |
| WT 2030_2808 | S2030A | 1.413E-01 | 0.00833 | |
| WT 2030_2808 | S2030A_Iso | 8.379E-01 | 0.01000 | |
| WT 2030_2808 | S2808A | 1.985E-01 | 0.01250 | |
| WT 2030_2808 | S2808A_Iso | 8.291E-10 | 0.00096 | REJECT |
| WT 2030_2808_Iso | S2030A | 1.749E-15 | 0.00076 | REJECT |
| WT 2030_2808_Iso | S2030A_Iso | 1.010E-12 | 0.00082 | REJECT |
| WT 2030_2808_Iso | S2808A | 5.083E-07 | 0.00106 | REJECT |
| WT 2030_2808_Iso | S2808A_Iso | 5.933E-01 | 0.01667 | |
| S2030A | S2030A_Iso | 1.488E-01 | 0.02500 | |
| S2030A | S2808A | 4.505E-03 | 0.00161 | |
| S2030A | S2808A_Iso | 2.115E-14 | 0.00077 | REJECT |
| S2030A_Iso | S2808A | 9.903E-02 | 0.05000 | |
| S2030A_Iso | S2808A_Iso | 7.368E-12 | 0.00085 | REJECT |
| S2808A | S2808A_Iso | 5.321E-07 | 0.00109 | REJECT |

### Supplemental Figure 1

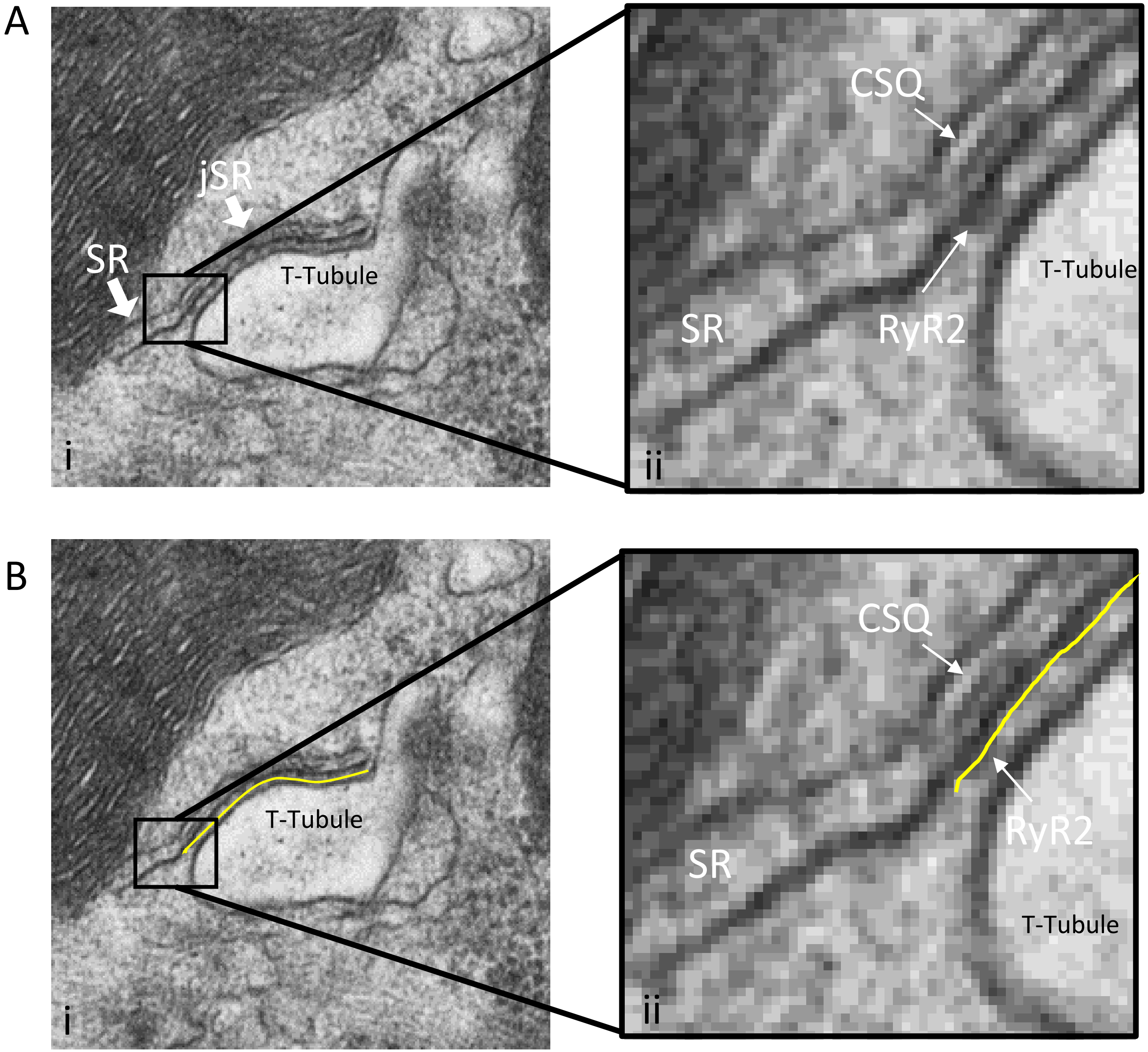


Supplementary Figure 1
Measurement of the Dyad Length.
The lengths of the dyads, regions of close apposition between junctional SR membranes with either a t-tubule or surface membrane, were measured using Fiji (Image j). Ai, jSR adjacent to a t-tubule. Extended SR is also visible in this image (SR), and has neither CSQ nor RyR2. Aii, Insets of the indicated regions in Ai are magnified 7×. Bi, Yellow line displays the area measured as jSR. Bii inset of the same region as Aii with the yellow line showing the exact area measured as the length of the dyad in all 2D TEM images.

### Supplemental Figure 2

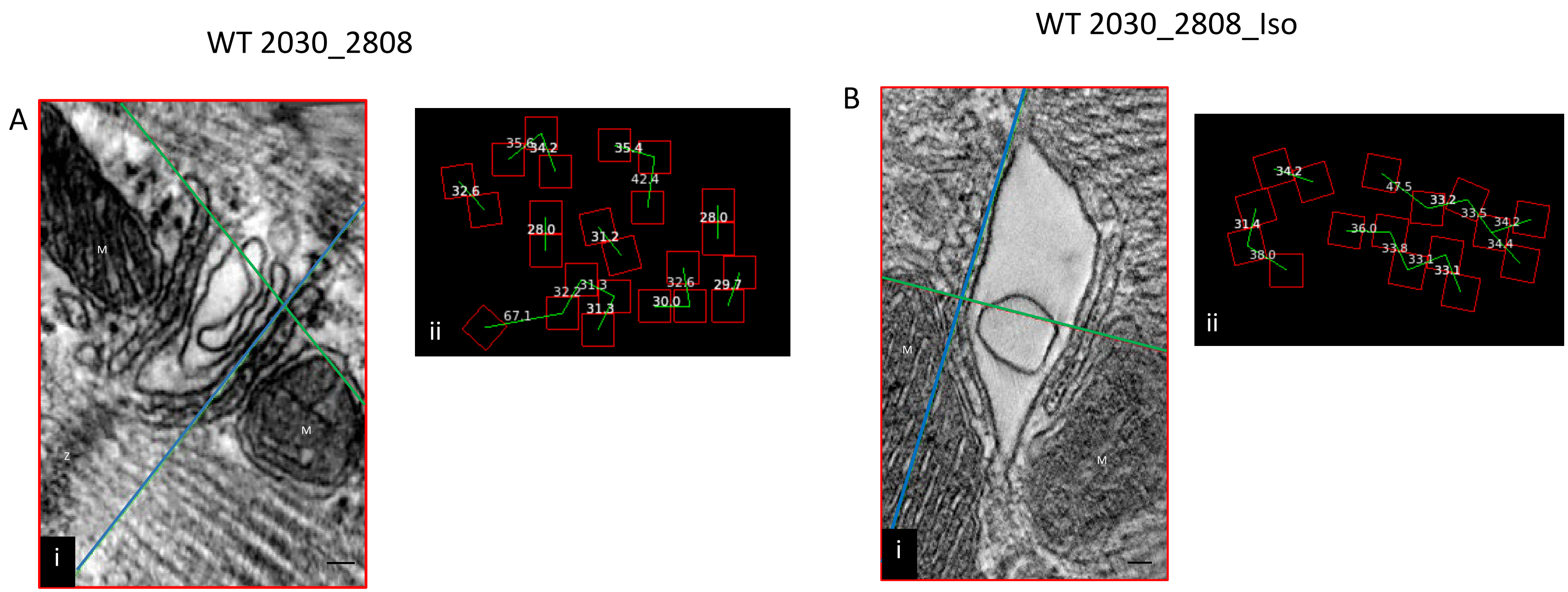


Figure Ai is a single image plane extracted from the dual-tilt tomogram of a mouse left ventricular myocyte fixed *in situ*. Different orthogonal planes from within the volume of the tomogram are indicated by different colors; XY in red, YZ in green and XZ in blue. The intersection point of all three planes lies within a single ryanodine receptor and the XZ plane (blue line) was positioned to parallel, as nearly as possible, the jSR membrane, but to be within the cleft and to bisect the ryanodine receptors. Scale bar is 30 nm. Figure Aii shows an enlarged view of the tetramer distribution (red boxes, 27 $nm^2$), and their NND. Figure B represents the same data for mice that had been treated with 300 nmol/L Iso for two minutes. M- mitochondrion, Z – Z line.

Supplementary Figures 2-7 display images for WT2814, S808A, S2030A, S2814A and S2814D, respectively.

### Supplemental Figure 3

WT 2814

A

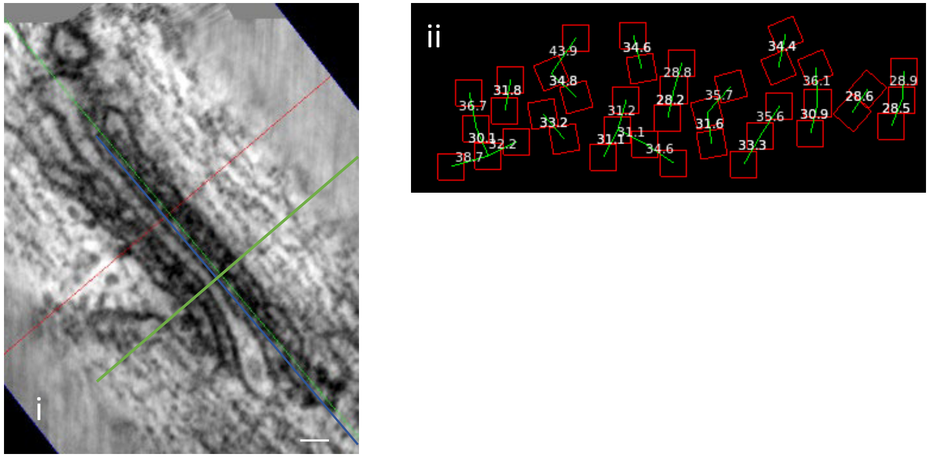


WT 2814_Iso

B

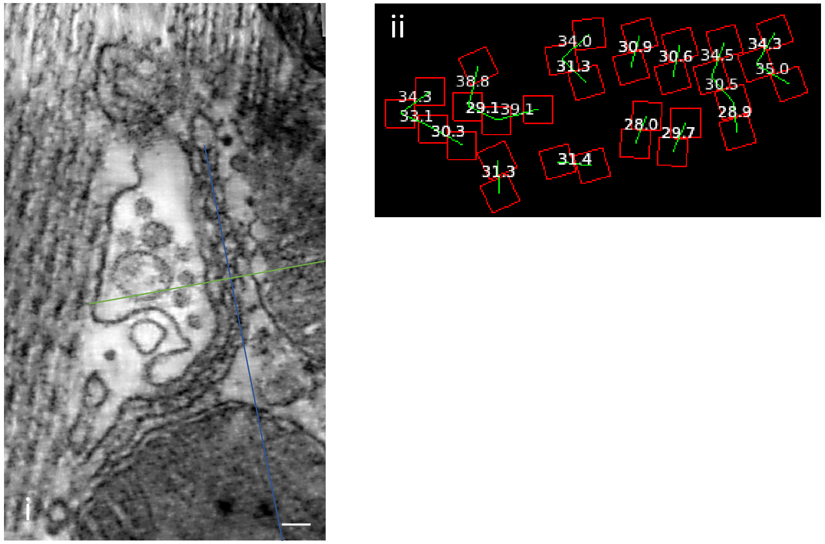

### Supplemental Figure 4

S2808A

A

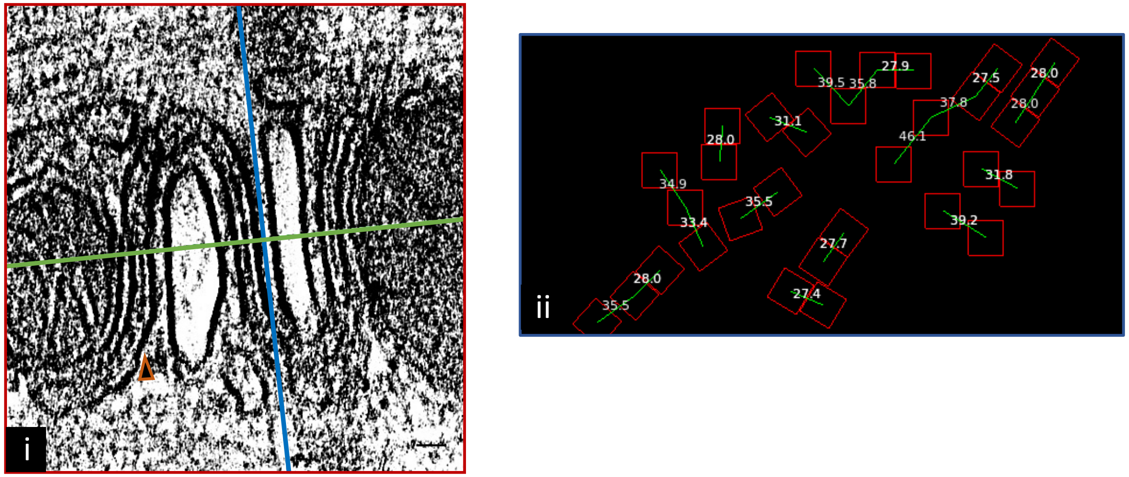


S2808A_Iso

B

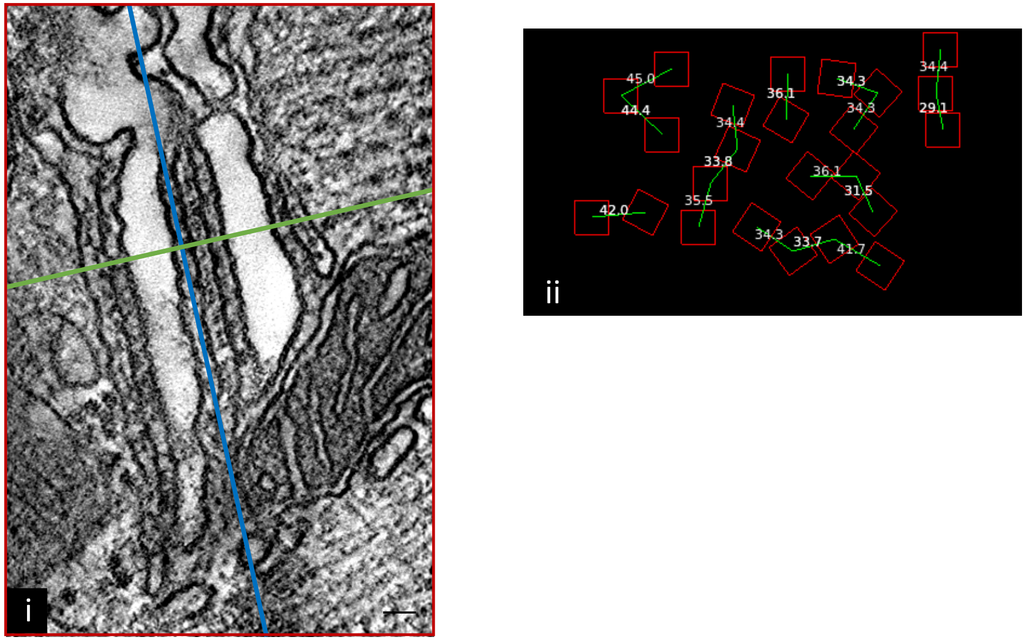

### Supplemental Figure 5

S2030A

A

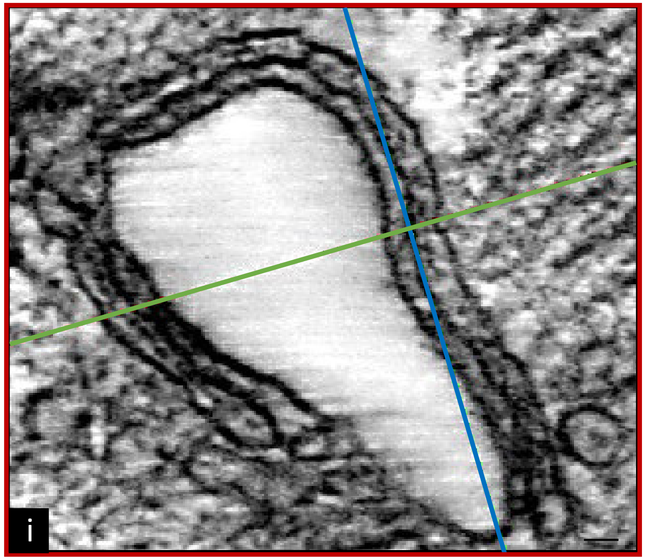


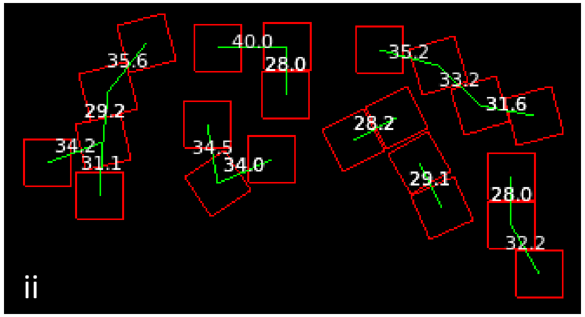


S2030A + ISO

B

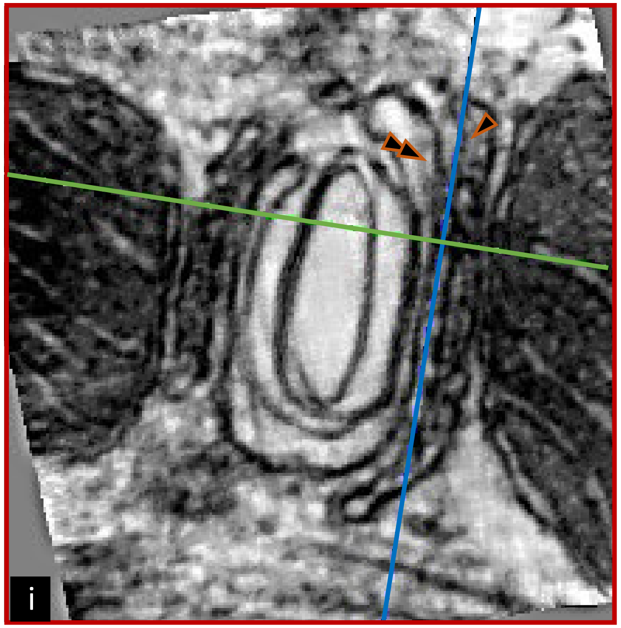


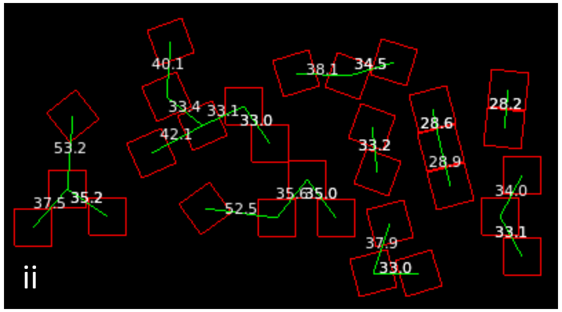

### Supplemental Figure 6

S2814A

A

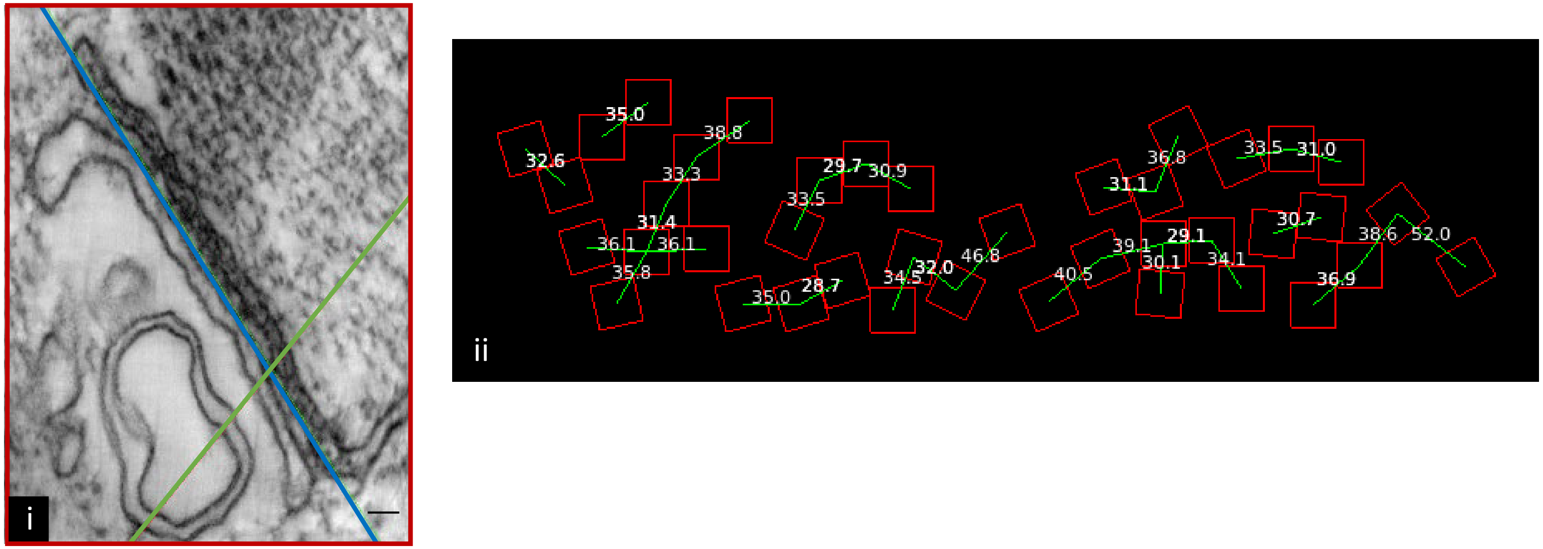


S2814A_Iso

B

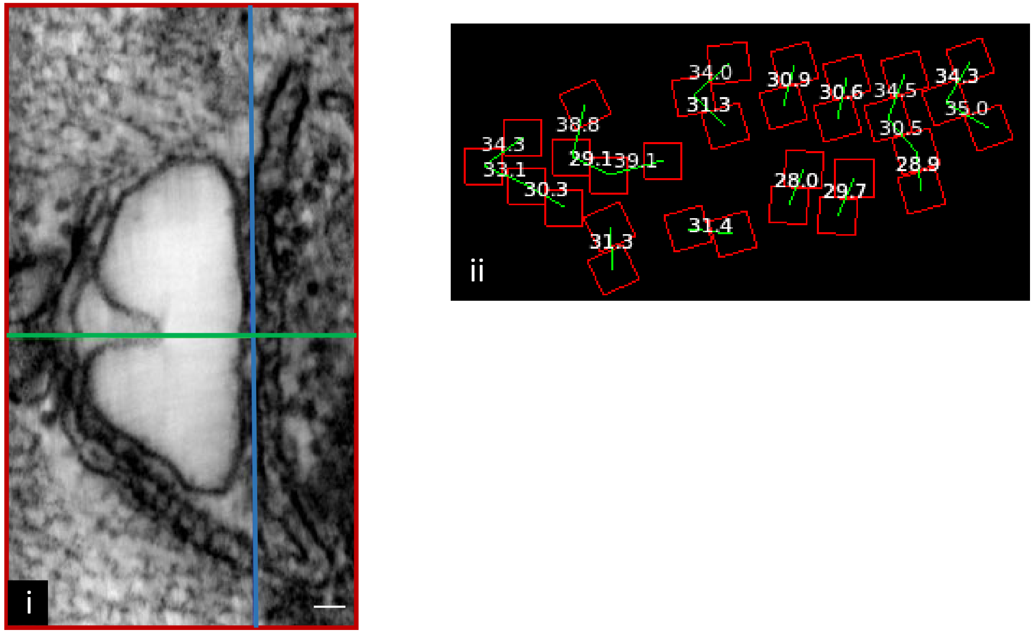

### Supplemental Figure 7

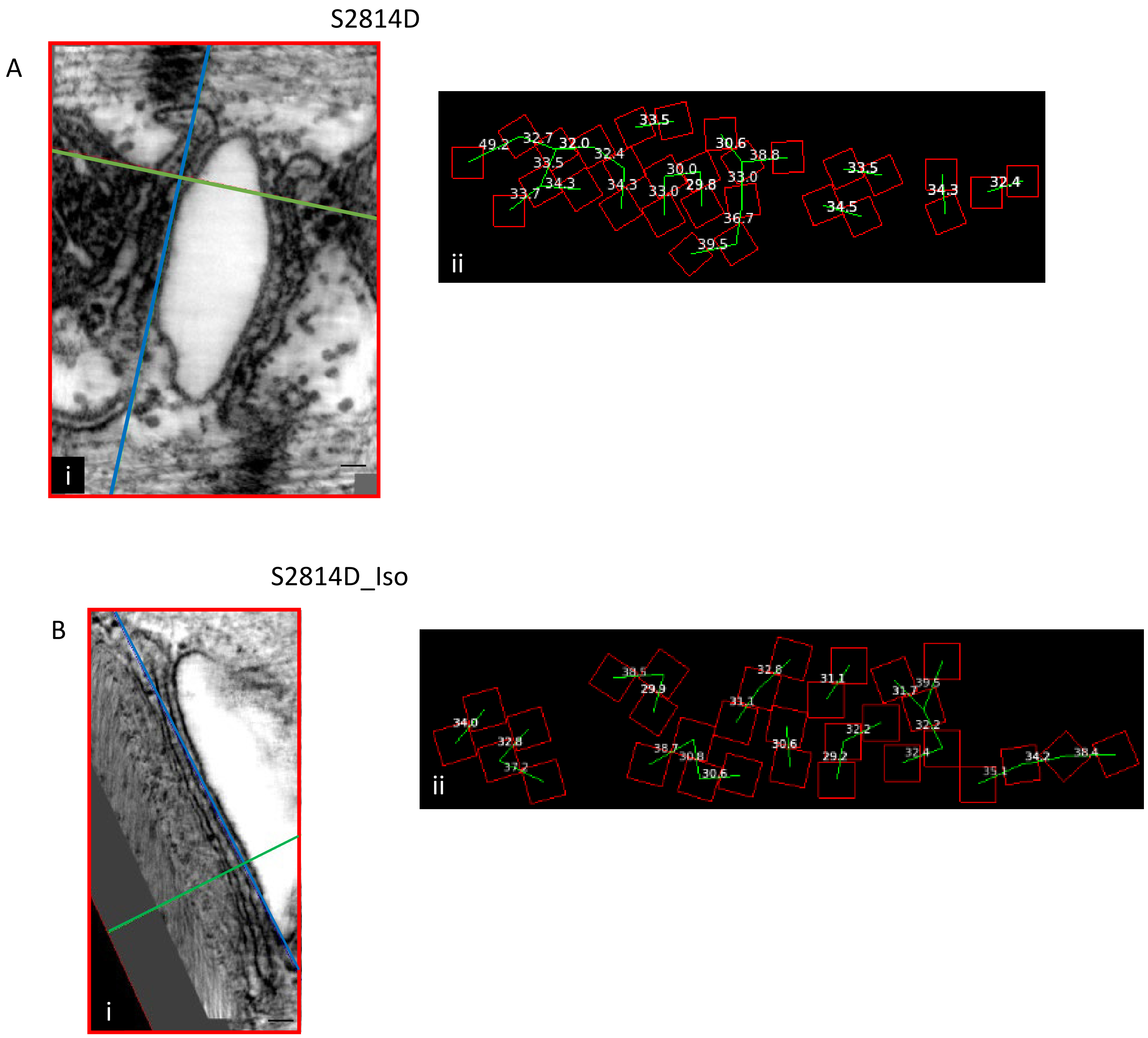
